# A schema for digitized surface swab site metadata in open-source DNA sequence databases

**DOI:** 10.1101/2022.12.15.520583

**Authors:** Barry Feng, Devin Daeschel, Damion Dooley, Emma Griffiths, Marc Allard, Ruth Timme, Yi Chen, Abigail B. Snyder

**Author notes:** Corresponding Author: Abigail Snyder.

## Abstract

Large, open-source DNA sequence databases have been generated, in part, through the collection of microbial pathogens from swabbing surfaces in built environments. Analyzing these data in aggregate through public health surveillance requires digitization of the complex, domain-specific metadata associated with swab site locations. However, the swab site location information is currently collected in a single, free-text “isolation source” field promoting generation of poorly detailed descriptions with varying word order, granularity, and linguistic errors, making automation difficult and reducing machine-actionability. We assessed 1,498 free-text swab site descriptions generated during routine foodborne pathogen surveillance. The lexicon of free-text metadata was evaluated to determine the informational facets and quantity of unique terms used by data collectors. Open Biological Ontologies (OBO) foundry libraries were used to develop hierarchical vocabularies connected with logical relationships to describe swab site locations. Five informational facets described by 338 unique terms were identified via content analysis. Term hierarchy facets were developed as were statements (called axioms) about how entities within these five domains were related. The schema developed through this study has been integrated into a publicly available pathogen metadata standard, facilitating ongoing surveillance and investigations. The One Health Enteric Package is available at NCBI BioSample beginning in 2022. Collective use of metadata standards increases the interoperability of DNA sequence databases, enabling large-scale approaches to data sharing, artificial intelligence, and big-data solutions to food safety.

**IMPORTANCE:** Regular analysis of whole genome sequence data in collections such as NCBI’s Pathogen Detection Database is used by many public health organizations to detect outbreaks of infectious disease. However, isolate metadata in these databases are often incomplete and poor quality. These complex raw metadata must often be re-organized and manually formatted for use in aggregate analysis. These processes are inefficient and time-consuming, increasing the interpretative labor needed by public health groups to extract actionable information. Future use of open genomic epidemiology networks will be supported through the development of an internationally applicable vocabulary system to describe swab site locations.

## INTRODUCTION

Modern surveillance of foodborne pathogens is reliant on large, open-source DNA sequence databases. Examples of laboratory networks dedicated to these surveillance programs include GenomeTrakr and PulseNet. Regular analysis of whole genome sequence (WGS) data in collections such as NCBI’s Pathogen Detection Database is used by many public health organizations to detect outbreaks of infectious disease [1]. In these analyses, the identification of pathogens from the environment that cluster with pathogens from sickened individuals informs epidemiological investigations [4]. Subsequently, the metadata associated with environmental pathogens directly support identification of point sources (Fig. 1). Regulatory environmental monitoring activities involve the collection of surface swabs within built environments followed by evaluation of infectious pathogens collected on the swabs [2,3]. The metadata that describe these swab site locations provides context for pathogen isolation source and must provide sufficient detail to be actionable [4–6]. In addition to contextualizing data from individual samples, environmental monitoring metadata enables large collections of genomic data to be shared and integrated through digital networks [7]. Realization of such open genomic epidemiology networks has been hindered by the absence of an internationally applicable vocabulary system to describe swab site locations.

**Figure 1.**
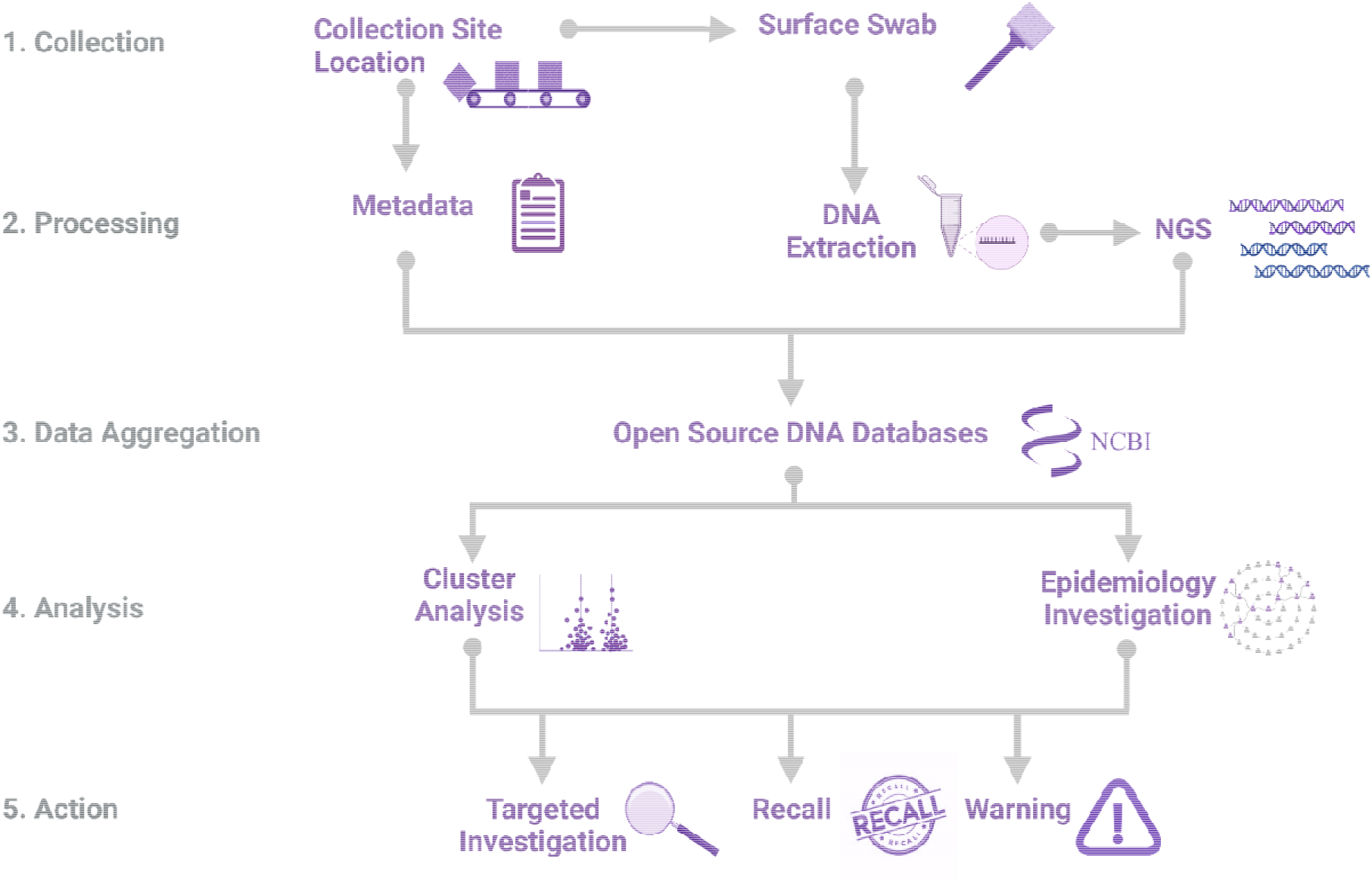
An overview showing the investigation of infectious disease outbreaks using WGS and associated source metadata. Abbreviations: WGS, whole genome sequencing.

Despite its importance, isolate metadata in large, open-source DNA sequence databases are often incomplete and poor quality. Metadata include general attributes (date, geographic location) as well as domain-specific attributes (swab site location, collection method) [4,9,10]. Descriptions of swab site locations within built environments are particularly challenging to standardize due to the complexity and variation of the necessary information and, consequently, are currently reported by individual data collectors as unstructured, free-text responses. These complex raw metadata must often be re-organized and manually formatted into a uniform pattern to be usable in aggregate analysis. Primary data collectors must sometimes be contacted during outbreak investigations for additional information. These processes are inefficient and time-consuming, increasing the interpretative labor needed by public health groups to extract actionable information to address urgent public health needs [4]. By contrast, improved metadata standards support FAIR principles of data management, enabling increased Findability, Accessibility, Interoperability, and Reusability of large, open-source DNA sequence data [10].

The goal of this study was to use a collection of unstructured, free-text swab site location descriptions to inform the development of a schema to structure and standardize swab site location metadata. Such a system will be integrated within broader minimum metadata standards for the built environment and used by data collectors within public health groups [7,8,11]. To accomplish this goal, Open Biological Ontologies (OBO) Foundry Principles [12] were used to develop a schema that defined (1) informational facets, (2) ontologized terms, and (3) statements (called axioms) about how entities within informational facets were related, to better structure and standardize descriptions of environmental monitoring swab site locations. We then applied this schema in a use-case of *Listeria monocytogenes* from different food production environments. Collectively, this analysis allowed us to identify gaps in existing ontologies, such as the lack of terms for industrial equipment, and create resources appropriate for our use-case and applicable for re-use in the analysis of other types of datasets to better harmonize and integrate public health food safety research.

## METHODS

### Collection and evaluation of unstructured, free-text metadata

We studied 1,498 swabs site descriptions generated during routine food safety surveillance and investigation activities by the U.S. Food and Drug Administration (FDA). All information specific to the facility was anonymized. These records were from a total of nine facilities on 22 different collection dates selected through convenience sampling. Sites included one dairy, four produce, one seafood, and one mixed facility. All text from the swab site metadata were extracted using the text mining tool (tm) v0.7-8 in R version 4.1.2 [13]. Free-text responses were anatomized by categorizing the components of each description into different data facets. In addition to extracting word-frequency counts, content analysis was used to analyze themes and patterns of the free-text responses based on explicit rules [14]. Briefly, emergent coding was performed by two researchers who independently reviewed the free-text metadata and identified a set of features that formed an initial checklist. The researchers then used the consolidated checklist to independently code features in free-text metadata. An example of how different concepts were formalized as informational fields is illustrated in Supplemental Fig. S1. After independently coding features, the agreement between the researchers was >95%. In cases of disagreement, the coding was discussed and a consensus on assignment identified.

### Proposing new terms to OBO

Each unique free-text term was queried in the ontology lookup service (OLS) ONTOBEE [13,15]. Terms that were not found in ONTOBEE were proposed as new terms within existing Open Biological Ontologies (OBO). Proposed new terms were each linked to an existing OBO term which served as the parent class representing the broader hierarchical category where the new term would be categorized. For example, the term *rusted* was linked to *texture* in the Phenotype and Trait Ontology (PATO). In addition, the textual definition of the proposing term was generated following the Aristotelian format [16]. The parent class and textual definition of terms were documented in a ROBOT template as shown in Supplemental Table S1. A summary of the information that must be documented within ROBOT templates to propose new OBO terms can be found in Supplemental Table S2. The ROBOT files were submitted to the relevant ontologies through their GitHub page. Following the submission, ontology curators initiated a quality check (QC) procedure to ensure that the semantics of the proposed terms were compatible with those of the existing terms. Terms passing the QC check were assigned an ID number and were integrated within the ontologies. The larger framework connecting these terms, described in detail within Results, utilized logical relationships from the Relation Ontology (RO) as summarized in Supplemental Table S3.

### Statistical analysis

A rarefaction curve was generated using the vegan package v2.5-7 with a subset of 20 unique descriptions to evaluate the diversity of terms collected from the free-text swab site metadata [17]. The swab site locations within our collection where *L. monocytogenes* was detected were analyzed as an example use-case for our schema. This data set is publicly available in Supplemental Table S2 and the *L. monocytogenes* genomic data is available on NCBI through the identified SRR numbers. A chi-square test of association with a Bonferroni correction was performed in R v4.1.2 to identify structure locations with significantly higher incidence of *L. monocytogenes*.

## RESULT and DISCUSSION

### Lexicon of unstructured, free-text metadata

Here we present an assessment of the lexicon typically used by data collectors in unstructured, free-text descriptions of swab site locations. By anatomizing these responses (breaking down descriptions into discrete concepts), we identified common language structures, as well as recurrent issues. These findings informed the development of the standardized schema described below. Within the free-text responses, the frequent use of synonyms and presence of occasional typos complicated machine readability in aggregate analysis. We have used a conveyor belt as an illustrative example in this article because it is a common and complex structure often associated with harborage of *L. monocytogenes* through surface swabbing. In free-text responses, data collectors variously referred to this structure as a *conveyor, conveyor system, processing line, conveyor belt, belt*, or simply by the brand name of the equipment, and a common misspelling was *conveyer*. Moreover, swab site descriptions were not single terms (eg., *conveyor*), but rather short descriptive statements such as *leg of the conveyor with rusted hole* which contained several different facets of information. This complexity contributes to the challenge of machine interpretation of swab site location descriptions. In addition to the potential complexity, free-text statements were also often incomplete or imprecise. For example, in the description *condensation coming down next to conveyor* it is unclear what surface structure was swabbed, beyond that it has condensation on it, which complicates even human interpretation.

Through assessment of term diversity used in free-text descriptions, we identified 338 unique terms used within 1,498 swabs site descriptions. The majority of these terms (n=253) were used to describe the structure and the sub-part of the structure that was swabbed. A rarefaction curve illustrating the increase in the number of unique structure terms as a function of the number of the total swab site descriptions (Fig. 2A) revealed a high level of term diversity used to describe surface structures [18,19]. Significant term diversity represents a challenge in the standardization of metadata format and signals an ongoing need for management and curation,[20].[21,22]. As a consequence of this finding, adoption of controlled terms, such as in a community-supported ontology, may be warranted as ontology provides a clear definition and promotes consistent application across data collectors,[23],[24].

**Figure 2.**
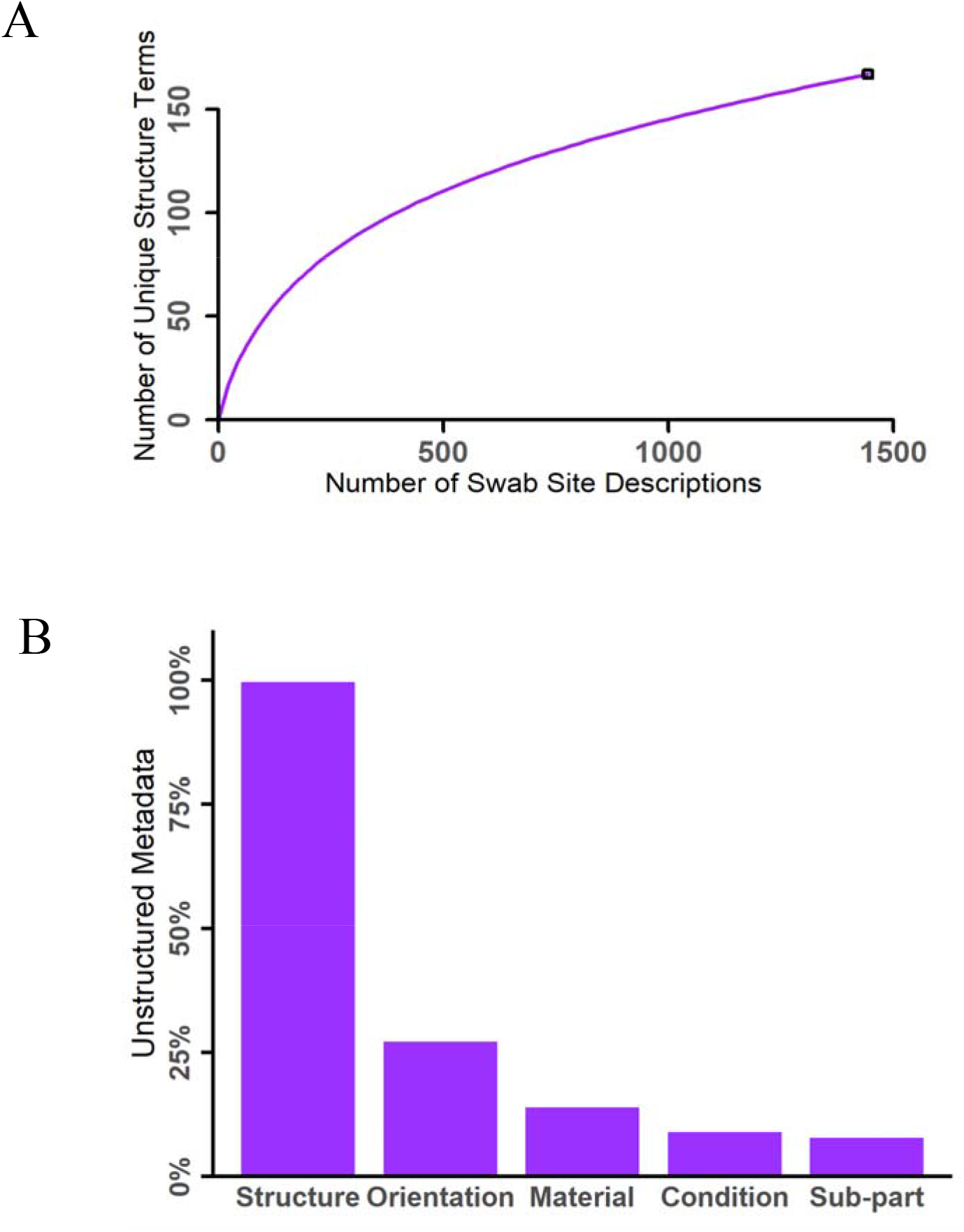
Analysis of free-text swab site location descriptions revealed (A) a high level of term diversity and (B) a low degree of completeness across information facets typically collected.

While the majority of unique terms used in free-text responses referred to the structure, several other informational facets describing important aspects of the surface environment were commonly included. In total, we identified five unique informational facets: 1) the structure being swabbed, 2) the sub-part of the structure, 3) the material from which the swab surface is composed, 4) the condition of the surface, and 5) the orientation of the swab site location on the structure. As an example, *underside of the cracked plastic belt of the conveyor* includes all five information facets. An assessment of how consistently each of these informational facets were addressed across unstructured, free-text responses revealed that while the structure was defined in nearly all metadata (99.6%), the remaining informational facets were addressed far less frequently (Fig. 2B). This suggests that the completeness of swab site descriptions could be improved by prompting data collectors for specific informational facets. By contrast, metadata dumped into a single “isolation source” field triggers the collection of short, less detailed description where word order, granularity, spelling, and the use of synonyms or abbreviations vary. In epidemiological investigations, the analysis and interpretation of the swab site information impacts the speed and scope of public health response [4]. Incomplete or uninterpretable metadata increases the difficulty of identifying the origin of infectious pathogens, which complicates root cause analysis [25].

### Ontology reuse and axiom construction

Ontologies are one solution to the challenges associated with the free-text responses. By defining terms in a hierarchical structure, ontologies standardize the definition for each term. For instance, the term *conveyor system* falls under the parent category *system* (Fig. 3). This hierarchy defines how a conveyor belt is related to other types of manufacturing equipment, which is important for swab site descriptions that contain different levels of granularity. Comparison among metadata that vary in granularity is otherwise difficult without substantial text-mining and is currently limited by agency-specific classification schemes which are essentially flat lists. Because ontological terms are connected through logical reasoning, definitions for these terms are more explicit and encode knowledge within a specific domain [12,24].

**Figure 3.**
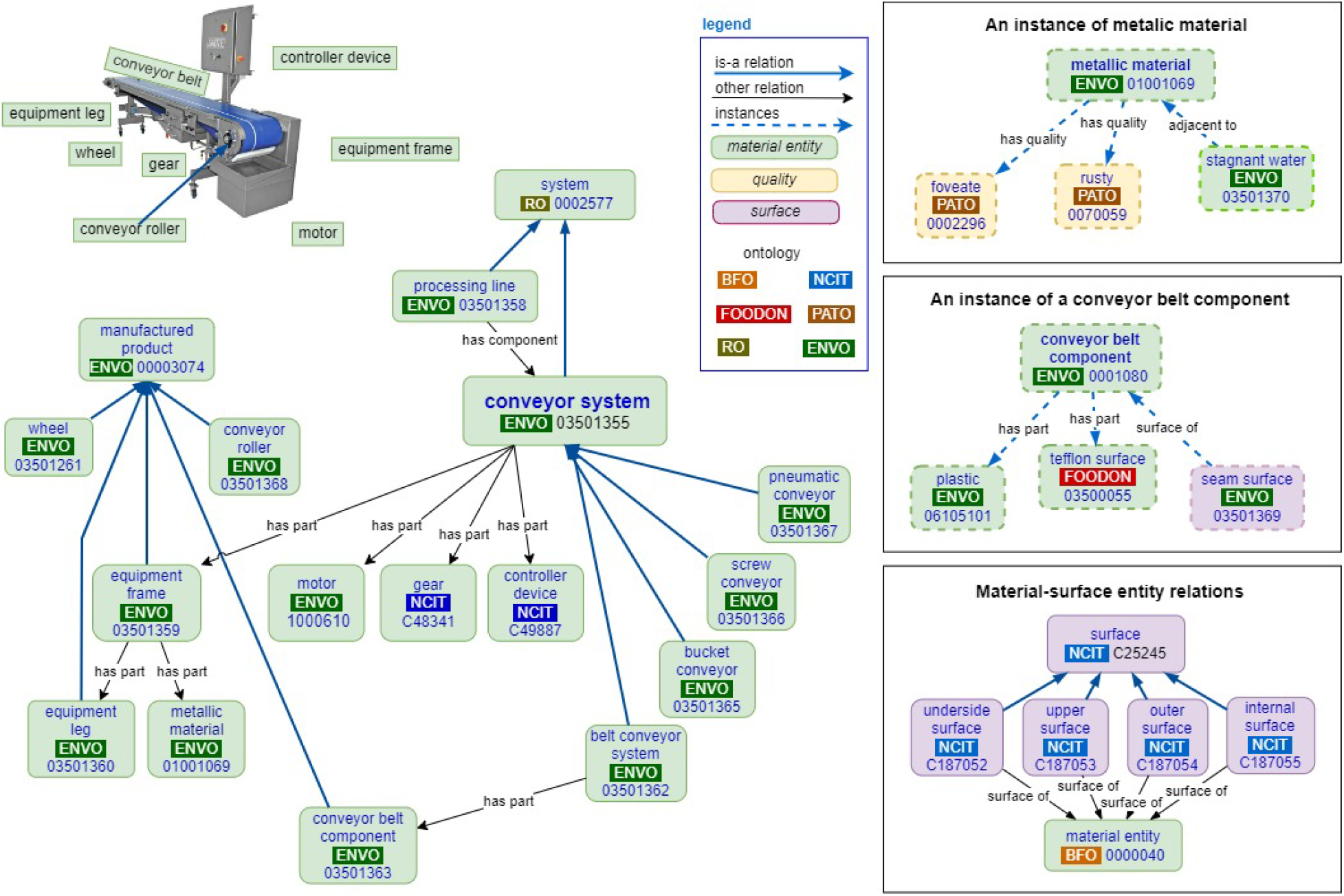
A semantic model illustrating the logical relationships among unique terms representing different informational facets used to describe swab site locations on a conveyor belt.

Most of the terms we identified in free-text responses were transformed from the Environment Ontology (ENVO) or the Phenotype and Trait Ontology (PATO). ENVO contains specialized terms for industrialized equipment and manufacturing applications, supporting informational facets related to the structure or the sub-part of the structure being swabbed [21]. PATO contains terms describing the characteristics of material things, supporting informational facets related to the type of surface material and its condition. Both are mature community-supported ontologies. Ontologies enable community consensus about classification schemes and the meaning of descriptors. Historically, metadata has not been documented using this vocabulary and terms used in free-text metadata are less nuanced. OBO Foundry principles encourage the re-use of terms from other ontologies wherever applicable [26,27]. This prevents redundant or conflicting efforts and maximizes the benefit from community coordinated efforts in maintenance of centralized registries. Use of third-party ontologies allows users to access and retrieve terms from specialist domain ontologies and new terms, when needed, can be proposed. For example, we proposed several new terms to ENVO such as *equipment leg* (ENVO: 03501360), *conveyor roller* (ENVO: 03501368) and to PATO including *rusted* (PATO: 0070059) which were not previously included, but which are relevant to our use case. Term proposal is a necessary and continuous effort to adoption of an ontology approach. For example, we found gaps in ENVO due to a lack of terms describing manufacturing equipment. Similarly, we found that terms in PATO centered biological systems which did not capture all nuances of the characteristics describing abiotic surfaces. Our term proposal focused on addressing these gaps, which will be an ongoing necessity for dedicated user groups.

Adoption of ontologies standardizes the use of terms by data collectors, enforcing accurate referencing. Terms are linked to OBO ontology identification numbers and object properties, thus avoiding the use of ambiguous free-text responses. Controlled definitions promote consistent application, in comparison to the examples of misspellings, synonyms, and colloquialisms often observed in unstructured, free-text responses [28]. Moreover, these definitions within term hierarchies are managed through cross-cultural and expert consensus in use-case domains, ensuring broader understanding and agreement [27,29]. We developed a semantic model (Fig. 3) to illustrate the conceptual relationship among the informational facets commonly used to describe swab site locations. Individual terms are connected with axioms which are statements about how entities within a domain are related [30–32]. These interconnections are held by the Relation Ontology (RO) and are represented by arrows shown in Fig. 3 [33]. For example, the “has part” RO statement establishes the *conveyor roller* as a subpart of the larger *conveyor system*. Whereas the “has quality” statement allows reference to material condition features through various terms including *pitted, rusted*, or having *standing water*, as relevant. It is notable in this example that some of the “has quality” characteristics can be described as *instances* and are therefore represented in a dashed line within this visualization (Fig. 3). The use of instances implies that these characteristics, such as *rusted*, apply to some but not all metallic materials. Ultimately, this framing supports the machine-readability of metadata by assigning ontology identification numbers that can be recognized digitally and support computer recognition of semantic relationships [34].

### Application in the “smarter” era of food safety

The schema described here has been implemented in the One Health Enteric Package, an expanded and standardized suite of metadata for genomic surveillance of enteric pathogens [35]. This package was developed by a U.S. interagency working group, GenFS [39] with the goal of expanding and standardizing metadata collected for sample types that span the One Health continuum: humans, animals, and environments including built environments [36]. Further, ontology-based packages like this make comparison across other ontology-based schemes developed by other agencies (with overlapping but slightly different scopes) more mappable and easier to compare. For example, the U.S. interagency One Health Enteric Package may be more readily compared with the One Health AMR standard being developed by a joint-agency Canadian initiative [37]. Complete and consistent metadata enhances the efficiency of epidemiological investigations, as evidenced by other groups who have successfully applied ontology-based approaches including NCBI and EMBL-EBI [27,38]. However, to our knowledge, this is the first schema that has been developed to capture machine-readable descriptions of swab site locations in built environments, building on the MixS minimum metadata standard from for built environments [7]. In application, data collectors can draw upon this schema to standardize vocabulary usage in their interest domain. However, this also requires buy-in from a broad base of data collectors who may need to expend more effort in the generation of standardized metadata compared to the level of effort currently needed to generate free-text responses. While this upfront effort ultimately benefits public health by expediting investigations and reducing interpretative labor [4], buy-in from data collectors can also be enhanced by reducing the barriers to adopting best practices. For example, application-based tools that increase the ease of metadata collection will increase buy-in (Fig. 4A). Training, technical support, and a shared conceptual understanding of the value of these upfront efforts are also important to adoption. These efforts are more generally a part of U.S. FDA’s framework outlined in the “New Era of Smarter Food Safety Blueprint” targeting tech-enabled traceability and adoption of smarter tools and approaches for prevention and outbreak response [39]. In that vein, broader application of frameworks that standardize metadata increase automation and machine-actionability.

**Figure 4.**
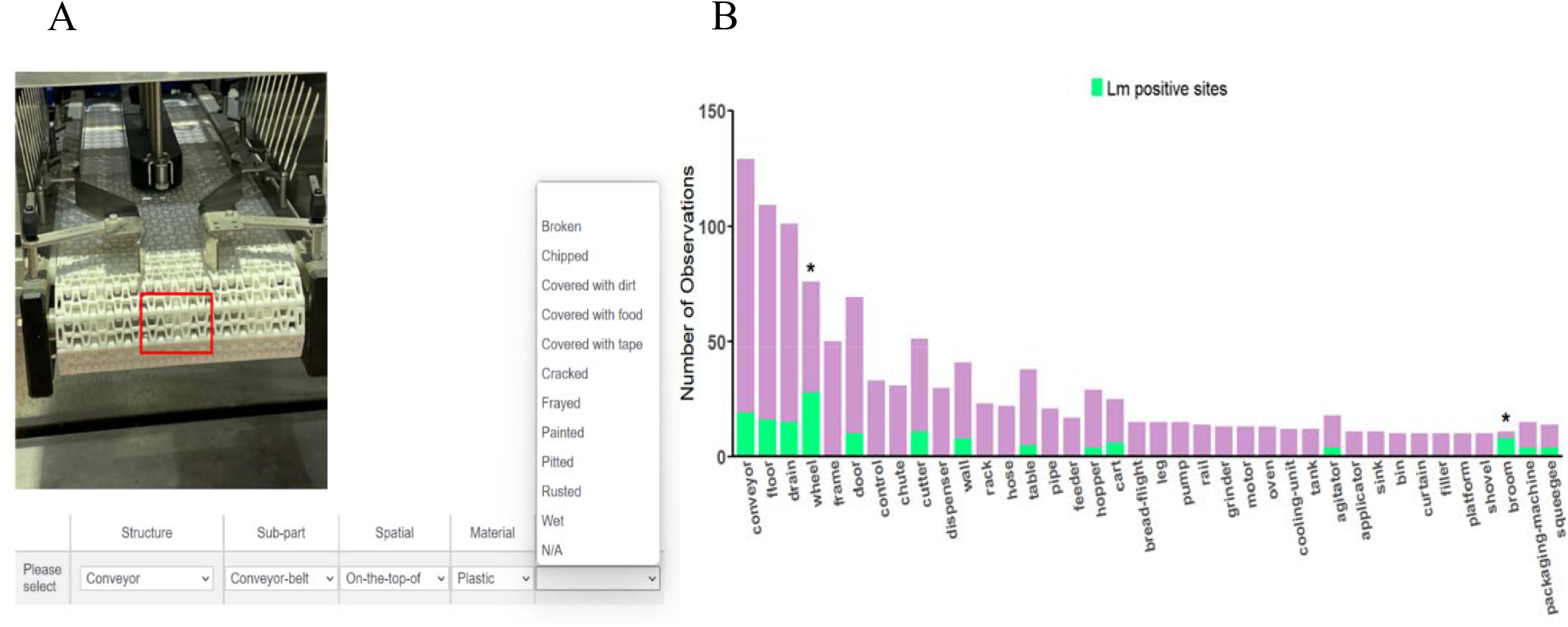
Standardized swab site location metadata supports public health initiatives. (A) Digital tools such as applications with selectable drop-down menus can assist data collectors in real-time data capture. (B) Machine readable metadata supports rapid assessments of sampling effort and pathogen detection. *Green shading corresponds to the number of swab sites that were positive for Listeria monocytogenes. (*) indicates structures where L. monocytogenes was significantly overrepresented*.

Digitized metadata also support more fundamental initiatives driven by machine learning, artificial intelligence, and big data approaches to food safety, beyond simply facilitating outbreak investigations. The internet of things (IoT) requires machine readable metadata and offers interconnection across people (eg, public health agencies, industry, consumers) and machines (eg, automated notification systems, open-source DNA sequence repositories, high-throughput bioinformatic pipelines) [27]. As a brief example, digitized metadata can enable quantitative evaluation of the surveillance efforts themselves. Decisions regarding which locations to sample are largely made by individuals, but machine-readable metadata allows evaluation of those selections in aggregate. From our data set we could easily identify what structures were most often selected for sampling once the metadata had been standardized and digitized (Fig. 4B). We could then assess the *L. monocytogenes* positivity rate among the most common swab site structures. Within this data set, wheels (28/128) and brooms (8/11) had significantly higher proportions of samples that were positive for *L. monocytogenes* (P<0.05) compared to other commonly selected locations. Although this example alone does not reflect a sufficient and balanced sampling on which to base policy, it illustrates how this type of analysis can drive improvement in targeting swab site locations as agencies iterate on their previous findings to identify high risk locations with increasing specificity. By contrast, analyzing these questions from free-text metadata would be prohibitively time consuming as NCBI’s Pathogen Detection database currently contains over 50,000 *L. monocytogenes* entries. As the rate of WGS data collection has increased dramatically over the last decade, it is crucial to adopt practices that ensure complete and high-quality metadata as quickly as possible for current and future database management [40].

## CONCLUSION

Outbreak investigation relies on the contextualizing information describing the source of environmental pathogens to track the origin of outbreaks of infectious disease. This metadata should be interpretable by humans and machines, providing sufficient detail for analyzing the source of pathogens. Based on our assessment of previously generated free-text responses describing swab site locations within built environments, we have identified common issues and proposed a schema for prospective, standardized metadata collection. This schema capitalizes on the framework from existing ontologies and formalizes data capture in five major informational facets. The resources developed here are also compatible with FoodOn [27] and GenEpio [41] which facilitates the integration of genomic epidemiological data with food product data. The One Health Enteric Metadata Package, hosted at NCBI BioSample, includes this framework for samples collected during facility inspections, in effect, implementing the schema recommended here for national and international genomic surveillance of foodborne pathogens.

## Supporting information

ROBOT FILE

## FUNDING

This work was supported in part by the U.S. Department of Agriculture, National Institute of Food and Agriculture [Hatch Project 2020-21-192] to ABS.

### Conflict of interest

The author declares no conflict of interest

## ACKNOWLEDGEMENT

We want to thank Dr. Erika Mudrak of the Cornell Statistical Consulting Unit and Dr. James B. Pettengill of the U.S. FDA.

## Notes

### Competing Interest Statement

The authors have declared no competing interest.

