## Supplemental Figures for "A schema for digitized surface swab site metadata in open-source DNA sequence databases"

**A**

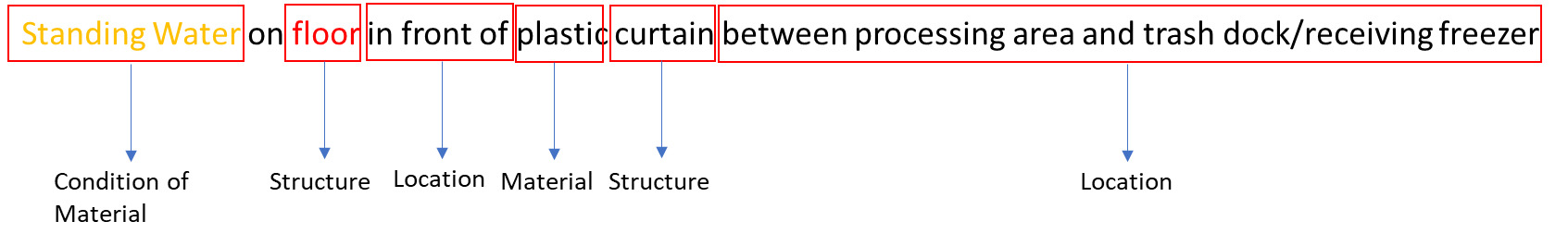

**B**

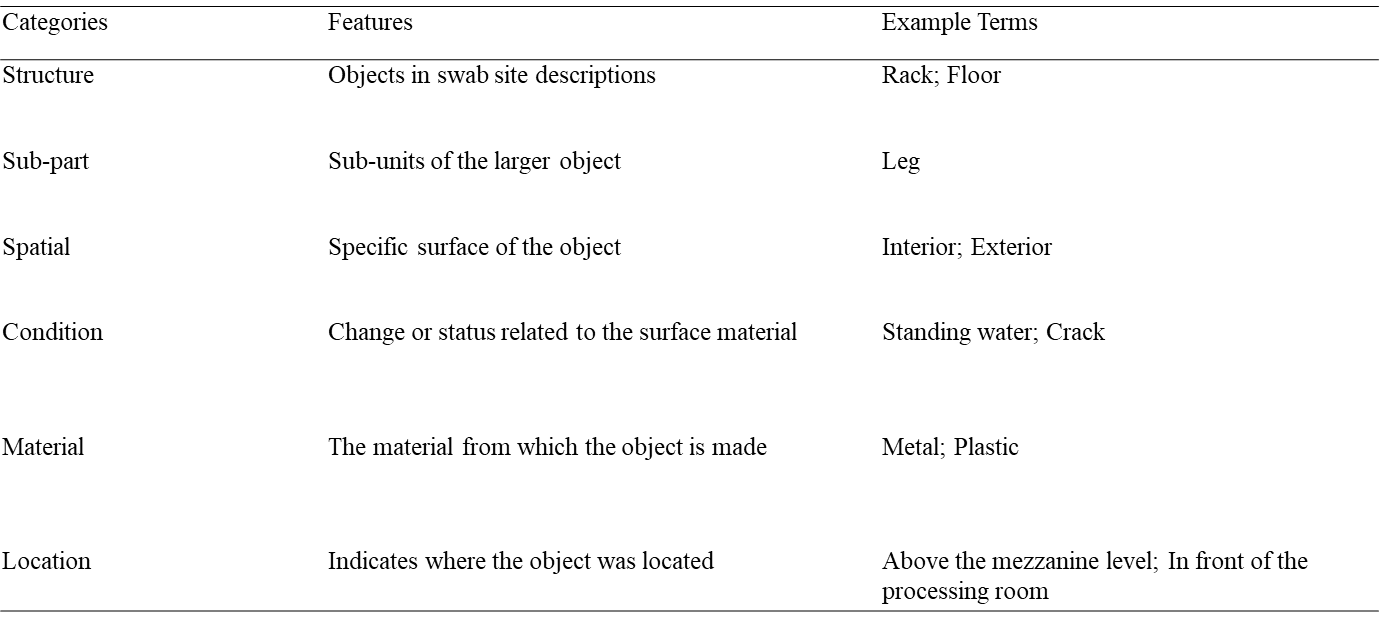

**C**

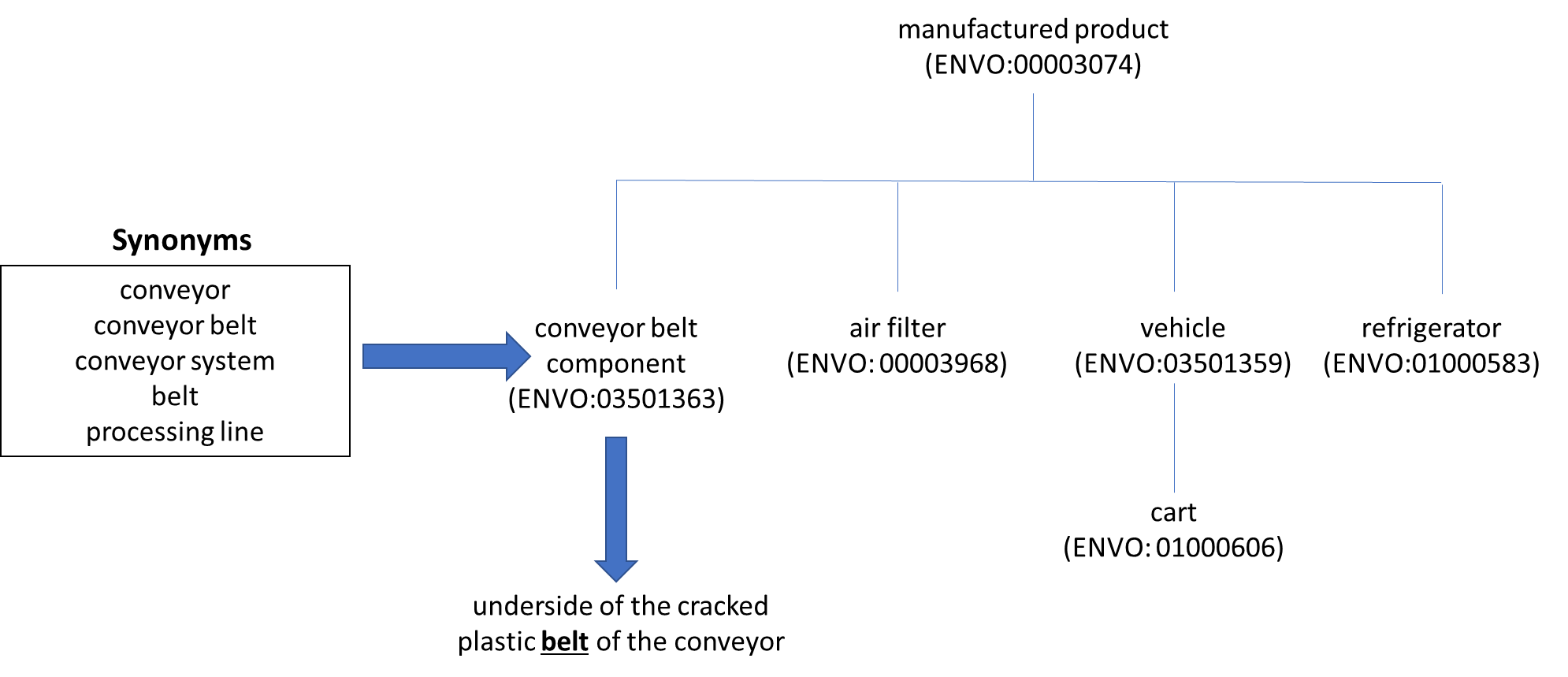

**Figure S1.** Metadata from unstructured swab site descriptions were categorized based on the following scheme. A) An example showing how an unstructured swab site description was anatomized based on this scheme. B) Descriptions and examples for the metadata categories in swab site descriptions. Out of these categories, structure, sub-part, orientation, material, and condition were identified as the five minimum required categories for metadata describing swab sites. Location terms were not included in the analysis. C) Illustration of how free-text terms were placed into hierarchical structures within ontologies.

**Table S1: A ROBOT template listing terms uploaded to the Environment Ontology (ENVO).**

**Table S2**: Information required for proposing terms to existing ontologies based on OBO principles. The example here is for the proposed entity “conveyor.”

| **Components** | **Description** | **Example** | **ROBOT label** |
| --- | --- | --- | --- |
| Ontology ID | An assigned ID number to represent the identity of the proposing term | ENVO:03501356 | ID |
| Label | The term that is proposing to the ontology | Conveyor system | A label |
| Parent class | The parent class of the proposing term | System | SC % |
| Definition | The definition of the ontology label in the form of [A X which has/is Y] | A system which is composed of one or more machines that can continuously transport material from one location to at least one other location. | AL IAO:0000115@en |
| Definition Cross Reference | Referenced source for the definition of the object | <http://vocab.getty.edu/page/aat/300024559> | >AI oboInOwl:hasDbXref SPLIT=\| |
| Comment | Additional information of the proposing term |  | AL rdfs:comment@en |
| Comment cross reference | Source of the additional information |  | >AI oboInOwl:hasDbXref SPLIT=\| |
| Editors note | Additional notes for ontology editors to better engineer to proposing term |  | AL IAO:0000116@en |
| Exact synonym | Other terms with the exact same meaning of the proposing term | conveyor equipment | AL oboInOwl:hasExactSynonym@en SPLIT=\| |
| Broad synonym | Synonyms with similar meaning as the proposing term but could also refer to other objects | conveyor | AL oboInOwl:hasBroadSynonym@en SPLIT=\| |
| Narrow synonym | Synonyms that refer to something more specific than the proposing term | food processing conveyor | AL oboInOwl:hasNarrowSynonym@en SPLIT=\| |
| Related synonym | Synonyms that could be narrow and broad sysnonyms |  | AL oboInOwl:hasRelatedSynonym@en SPLIT=\| |
| In subset | The community in which the proposing term is pertinent to | food safety | A oboInOwl:inSubset SPLIT=\| |
| Cross reference | Source providing the information of the proposing term |  | AI oboInOwl:hasDbXref SPLIT=\| |
| Subclass axiom | Other logical axioms helping to define the proposing term |  | SC % |
| Creation date | Date where the term was uploaded to ontologies | 2021-12-09T00:00:00Z | AT oboInOwl:creation_date^^xsd:dateTime |
| Created by | Creator's ORCID | https://orcid.org/0000-0002-2738-824X | AI oboInOwl:created_by SPLIT=\| |

**Table S3:** Terms in the Relation Ontology (RO) describing logical relations between entities relevant to swab site locations.

| **Logical Relationships** | **Description** |
| --- | --- |
| is_a | Represents the relationship in which an entity is the upper-class hierarchy of another object |
| has_component | Represents the relationship in which an entity is a sub-unit of another entity |
| material_surface_of | Represents the relationship in which an entity is a specific surface of another entity |
| has_part | Represents the relationship in which an entity is the material of another entity |
| has_quality | Represents the relationship in which an entity is the temporary state of a surface material |

**Table S4:** The SRR number of each *Listeria* strain listed in the metadata
